# The Higher Plant Plastid Complex I (NDH) is a Reversible Proton Pump that increases ATP production by Cyclic Electron Flow Around Photosystem I

**DOI:** 10.1101/049759

**Authors:** Deserah D. Strand, Nicholas Fisher, David M. Kramer

## Abstract

Cyclic electron flow around photosystem I (CEF) is critical for balancing the photosynthetic energy budget of the chloroplast, by generating ATP without net production of NADPH. We demonstrate that the chloroplast NADPH dehydrogenase complex (NDH), a homolog to respiratory Complex I, pumps approximately two protons from the chloroplast stroma to the lumen per electron transferred from ferredoxin to plastoquinone, effectively increasing the efficiency of ATP production via CEF by two-fold compared to CEF pathways involving non-proton-pumping plastoquinone reductases. Under certain physiological conditions, the coupling of proton and electron transfer reactions within NDH should enable a non-canonical mode of photosynthetic electron transfer, allowing electron transfer from plastoquinol to NADPH to be driven by the thylakoid proton motive force possibly helping to sense or remediate mismatches in the photosynthetic budget.

## Introduction

The canonical ‘Z-scheme’ model of linear electron flow (LEF, Figure S1A) uses a series of two photochemical reaction centers to extract electrons from water at photosystem II (PSII), and transfer them to NADPH through photosystem I (PSI) ^1^. The electron transfer reactions are coupled to proton transfer, generating an electrochemical gradient of protons—the proton motive force (Δp)—that drives the synthesis of ATP. Because of the strong coupling of electron and proton transfer reactions ^2,3^, LEF should produce a fixed (or rigid) ratio of ATP/NADPH [reviewed in ^4^]. In contrast, downstream metabolic reactions impose variable demands for ATP and NADPH, requiring dynamic adjustments of the photosynthetic energy budget to avoid “metabolic congestion” that can lead to buildup of products or depletion of substrates that can result in photodamage ^3,4^. Indeed, LEF alone should not be able to power the Calvin-Benson cycle. With one proton being deposited in the lumen during water oxidation and two being translocated by cytochrome *bf* complex through the Q-cycle, LEF should result in the translocation of 3 protons for each electron transferred to NADPH ^5–7^; with a H^+^/ATP ratio of 4.67 at the ATP synthase ^8^, LEF should produce 2.6 ATP for every 2 NADPH, a deficit of 0.4 ATP/2 NADPH compared to the ratio required to sustain the Calvin-Benson cycle.

There is substantial evidence that cyclic electron flow around photosystem I (CEF) plays an important role in balancing the ATP/NADPH energy budget ^4,9– 11^. The generally accepted model for CEF involves the transfer of highly reducing electrons from photoexcited PSI centers to plastoquinone (PQ) through a PQ reductase, resulting in the formation of plastoquinol (PQH_2_) with the uptake of protons from the chloroplast stroma. The PQH_2_ is then oxidized by the cytochrome *bf* complex and returned to the oxidizing site of PSI via plastocyanin or cytochrome *c*_6_.

There are several proposed CEF pathways in higher plant chloroplasts that differ at the level of the plastoquinone reductase [reviewed in ^12^]. One of these involves the thylakoid ferredoxin:plastoquinone oxidoreductase complex, for historical reasons called the NADPH dehydrogenase complex (NDH, Figure S1B, see Discussion), which is homologous to respiratory Complex I ^13^. Another pathway, termed the ferredoxin:quinone reductase (FQR, Figure S1C) is sensitive to antimycin A (AA), and proposed to involve a complex of the PGR5/PGRL1 proteins that is able to transfer electrons from ferredoxin (Fd) to PQ 12,14–17.

Both of these CEF routes are conserved across the flowering plants, even though they may appear redundant, i.e. both function in CEF by transferring electrons from ferredoxin to PQ ^15,16,18^. However, the PQ reductases in these two pathways are structurally distinct, possibly giving clues about their relative functions. Of particular interest is the homology of NDH to the bacterial or respiratory type I NADH:quinone reductases (Complex I) ^13,19,20^ that are known to couple reduction of quinones to the pumping of up to 2 protons for each electron transferred to quinone, efficiently generating Δp to drive ATP synthesis ^21,22^. In contrast, the FQR is likely to catalyze a simpler, more direct mode of PQ reduction ^14,16^ that is very unlikely to be coupled to proton translocation, much like the type II (non-proton-pumping) NAD(P)H:quinone oxidoreductases such as NDA2 and NDH2, as found in the mitochondrial respiratory chains of plants, certain fungi and protozoa. Non-protonomotive NDA/NDH2-type enzymes are also widespread in bacteria and the plastids of certain green algae ^23,24^.

If the NDH is functionally similar to Complex I, it could have the capability to use the energy liberated during PQ reduction to translocate protons across the thylakoid against the electrochemical gradient and thus increase the H^+^/e^-^ stoichiometry of CEF. Though proton-pumping activity for NDH has been proposed in previous work ^25,26^ it has not been experimentally demonstrated. Such activity would have a large impact on the energy balance of the chloroplast, by allowing for more efficient balancing of the ATP/NADPH budget with relatively low turnover rates. On the other hand, as discussed below, a high coupling ratio for proton-pumping to electron transfer could impose significant thermodynamic and kinetic limitations, or even allow plastoquinol to reduce NADP^+^. In this work, we used a series of complementary approaches to assess the possibility that NDH acts as a proton pump both *in vitro* and *in vivo*.

## Results

### Sequence conservation in NDH of residues essential for proton-pumping

Figure 1 shows a structural cartoon of *Arabidopsis thaliana* NDH indicating the likely organization and conservation of charged residues within the membrane domain, which are considered to be essential for proton translocation in respiratory Complex I. Complete sequence alignments for the membrane subunits depicted in Figure 1 for the respective NDH subunits from *A. thaliana*, *Spinacia oleracea* (spinach), and *Nicotiana tabacum* and Complex I from *Escherichia coli*, *Thermus thermophilus*, *Yarrowia lipolytica* (yeast), and *Bos taurus* are presented in Figure S2(A-E).

**Figure 1.**
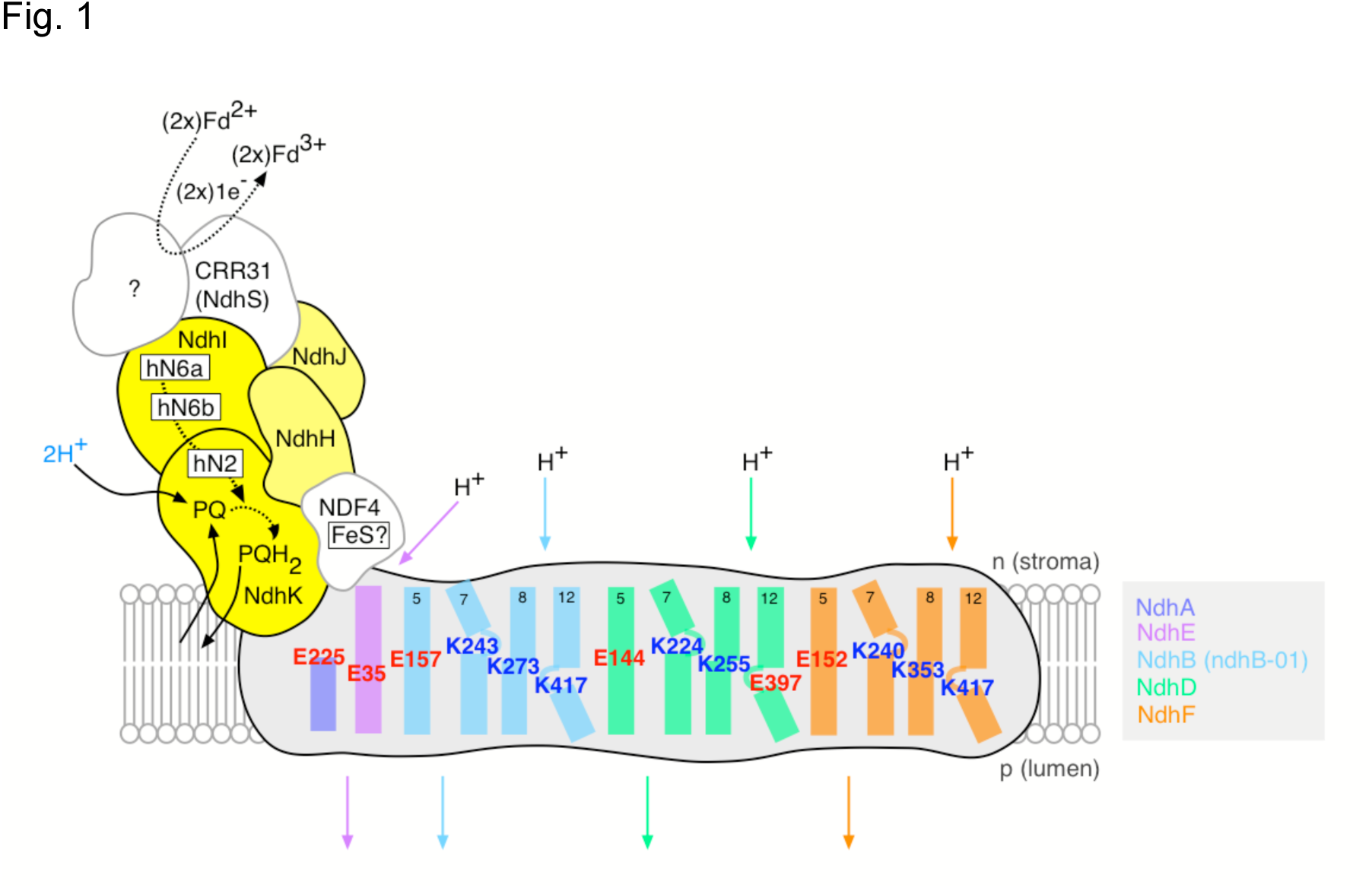
Structural cartoon of NDH showing conserved residues essential for proton-pumping within the membrane domain (grey) and likely organization of the electron donor domain (yellow). Subunits equivalent to the NADH-binding domain of respiratory Complex I (the ‘N-module’ in the terminology of Brandt ^63^) have not been identified in NDH. The (plasto)quinone-binding ‘Q-module’ is extant in NDH, and formed from NdhH,-I, -J and –K. Structural motifs for the Complex I iron-sulfur clusters N6a, N6b (NdhI) and N2 (NdhK) are conserved in NDH, although their presence has yet to be confirmed spectroscopically. As such, the ‘h’ prefix for these putative clusters in indicates their homology to Complex I. The subunits NDF4 and CRR31/NdhS are unique to NDH. NDF4 is likely to contain and iron-sulfur (FeS) cluster, and CRR31/NdhS has been proposed to be a constituent of the Fd binding site (although it may not be a redox-active participant) ^64,65^. Other stromal subunits that have been identified as NDH components but are unlikely to directly participate in the electron transfer pathway are not shown here ^66,67^. Highlighted charged residues within NdhB,-D, -E and –F in the membrane domain are conserved between NDH and Complex I, and are considered essential for proton translocation. Residues are numbered according to the *A. thaliana* sequence data. Helices are numbered according to their Complex I homologues. Predicted discontinuities in helices 7 and 12 of NdhB,-D and-F as observed in the atomic structure of *T. thermophilus* Complex I are shown. Note that the NdhB gene is duplicated in Arabidopsis (the sequences of B-01 and B-02 are identical). Complete predicted sequence alignments for the membrane domain subunits shown here are presented in Figure S2.

Four discrete proton channels, exhibiting highly characteristic sequence conservation, have been identified in the atomic structures of the membrane domains of *T. thermophilus* and *Y. lipolytica* Complex I ^27–29^. Three of these channels are contained within the ‘antiporter-like’ subunits equivalent to NdhB,-D, and-F, and the fourth from an association of subunits equivalent to NdhA,-C,-E and-G. This latter group also forms part of the quinone binding site within the enzyme. The lysine and glutamate residues located within the center of the lipid bilayer are key mechanistic features of these proton channels, and are conserved amongst Complex I and NDH (displayed in Figure 1). Mutagenesis of these residues has been shown to impair the proton-pumping capacity of *E. coli* Complex I [reviewed in ^22^]. The conservation of these intramembrane lysine and glutamate residues between Complex I and NDH provides circumstantial evidence for the protonmotivity of the latter enzyme, which was then investigated experimentally as described below.

### Proton-pumping activity of chloroplast NDH probed in vitro

*In vitro ATP generation in the dark via protonmotive NDH activity*. We developed an assay, described in Fig. 2A, designed to report ATP generation only in the presence of a proton-pumping, NDH-type plastoquinone reductase. ATP synthesis was monitored using firefly luciferase luminescence in spinach thylakoid preparations. Spinach was chosen as it provides a reliable (and abundant) source for the isolation of coupled thylakoids. To avoid interference from light-driven ATP production through photophosphorylation, experiments were conducted in strict darkness in the presence of dichloromethyl urea (DCMU), an inhibitor of PSII, and/or tridecyl stigmatellin (an inhibitor of the cytochrome *bf* complex, see below). The NDH reaction was initiated by addition of exogenous Fd, NADPH and the PQ analogue decyl-plastoquinone (dPQ). The suspension medium was well buffered so that the uptake of protons onto dPQ should not have affected the pH of the extra-lumenal space and thus Δp should only be produced by the transfer of protons in the thylakoid lumen. As shown in Figure 2B, addition of Fd or NADPH alone did not induce luminescence, but further addition of dPQ resulted in a sustained, reproducible increase in luminescence, reflecting the production of about 10 nmol ATP mg chlorophyll^−1^ min^−1^. The order of substrate addition was not critical, but that NADPH, Fd and dPQ were all required, implying that the oxidation of NADPH through Fd to dPQ was coupled to the synthesis of ATP (Figure S3). This result is consistent with the previously reported requirements of Fd for NDH activity ^18^.

**Figure 2.**
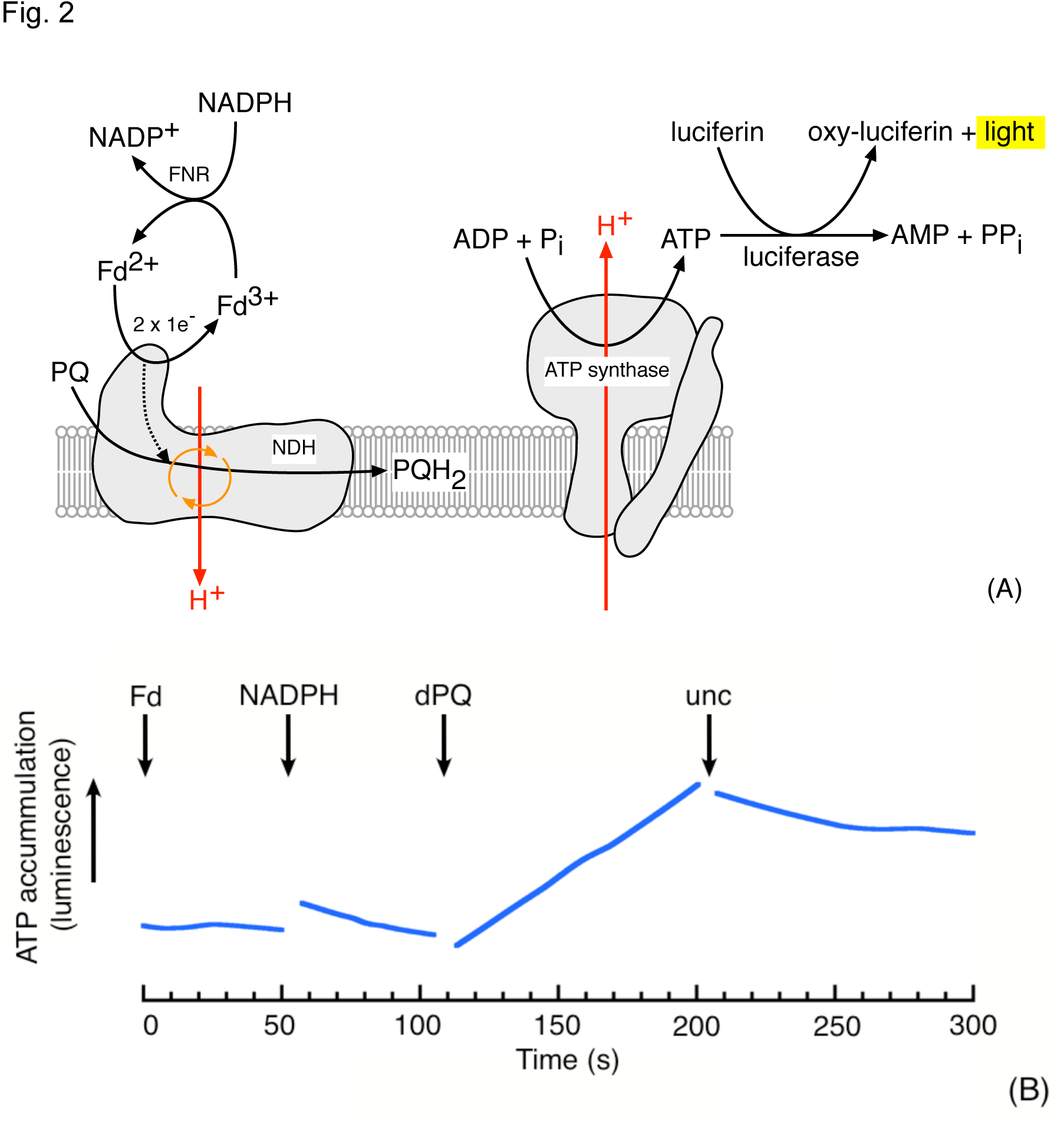
Evidence for involvement of a proton pump in CEF. (A) Cartoon of experimental rationale showing the coupling of proton-pumping NDH activity with ATP synthesis and subsequent detection by luciferase-mediated luminescence. A non-protonmotive NDH would be incapable of generating the proton gradient required for ATP synthesis. (B) ATP synthesis, monitored by luciferase luminescence, in DCMU-treated *S. oleracea* thylakoids (50 μg Chl/ml) in the dark after addition of 5 μM ferredoxin (Fd),100 μM NADPH and 50 μM decylplastoquinone (dPQ). The assay buffer consisted of 10 mM HEPES (pH 7.5), 10 mM KCl, 5 mM MglCl_2_, 2 mM ADP, 2 mM potassium phosphate, 2 mM DTT, 100 μM diadenosinepentaphosphate and 10 μM DCMU. Premixed ‘Enliten’ recombinant luciferase/luciferin reagent (used as supplied by Promega) was added to a final concentration of 8% (v/v). Addition of valinomycin and nigericin (10 μM each) is indicated by ‘unc’. Representative data (fitted by the locally weighted least squared error method) are shown; discontinuities in the data are due to the removal of mixing artifacts on substrate addition. The rate of ATP synthesis on dPQ addition was approximately 10 nmol ATP/mg Chl/min.

The observed ATP synthesis was dependent on the generation of Δp, as demonstrated by its abolition upon addition of uncouplers (valinomycin with nigericin, see arrow marked “unc” in Figure 2B). The ATP synthesis was insensitive to tridecyl stigmatellin (Figure S4), a potent inhibitor of the *bf* complex, indicating that the observed ATP synthesis also did not involve electron or proton translocation by the *bf* complex. Likewise, the activity was insensitive to oligomycin (Figure S4), a specific inhibitor of mitochondrial ATP synthase, indicating that the reaction was not catalyzed by contaminant mitochondrial respiration.

*Linkage of NDH activity to generation of* Δp *probed by post illumination fluorescence kinetics*. As a second, independent approach to determining the proton coupling of NDH, we assessed the effects of Δp on the reduction of the PQ pool using the “post-illumination fluorescence” signal, a transient rise in chlorophyll fluorescence observed in the dark after a sustained period of actinic illumination in leaves and chloroplast preparations. This fluorescence signal is generally considered to be related to the activity of NDH, and results from the transfer of electrons from stromal donors to the PQ pool, which equilibrate with the Q_A_ quinone molecule in photosystem II, resulting in elevated chlorophyll *a* fluorescence yield (Figure 3A) (see also ^13,19,30,31^).

**Figure 3.**
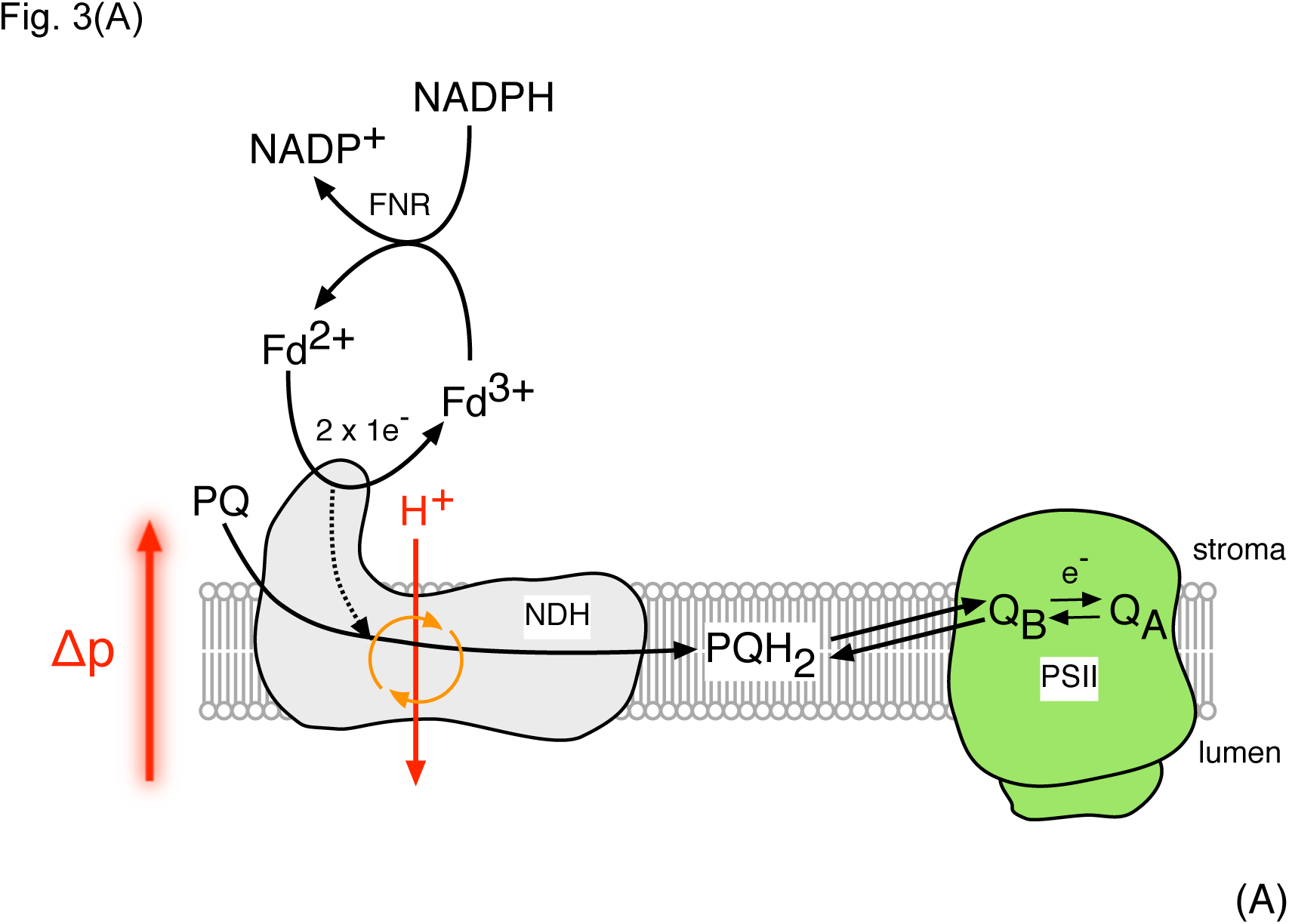
(A) Mechanism of association of NDH-mediated PQ pool reduction and the post illumination fluorescence rise. PSII, Q_A_ and Q_B_ represent photosystem II and the primary and second quinone electron acceptors in PSII respectively. As a protonmotive enzyme, NDH activity is subject to thermodynamic back pressure from the transthylakoid protonmotive force (Δp).

Figure 3B compares the kinetics of fluorescence yield changes in thylakoid preparations from *S. oleracea*, *Amaranthus hybridus*, *A. thaliana* (wildtype Col-0 and the NDH-knockout *ndhm* ^32^) using 100 μM NADPH as the electron donor in the presence of exogenous (spinach) Fd.

**Figure 3.**
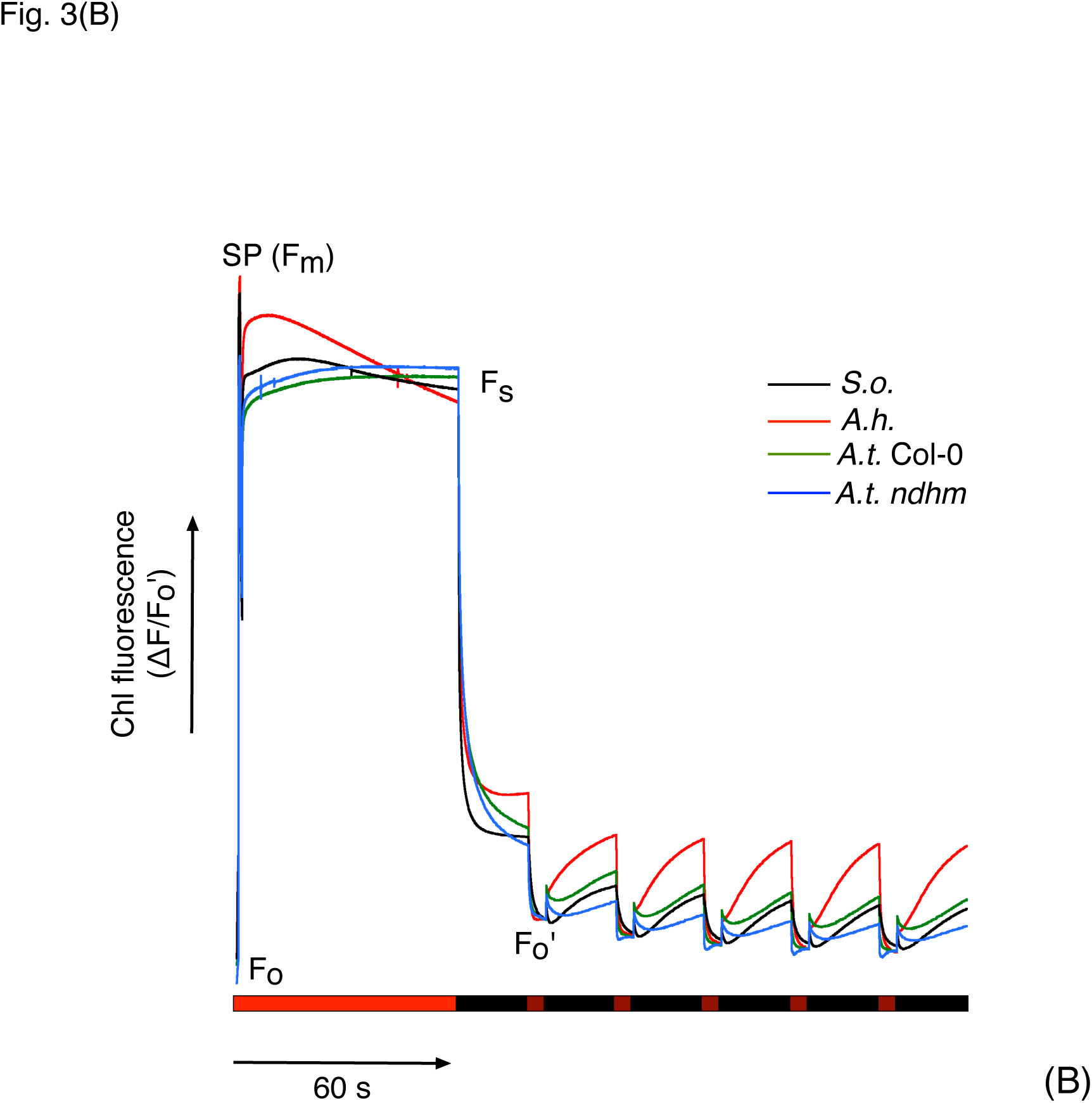
(B) The post illumination fluorescence rise in *Spinacia oleracea* (*S.o*), *Amaranthus hybridus* (*A.h*.), *Arabidopsis thaliana*- Columbia (*A.t*. Col-0) and *A.t. ndhM* thylakoid preparations. Assay conditions consisted of thylakoids suspended at 50 μg Chl/ml in 10 mM HEPES (pH 7.5), 10 mM KCl, 5 mM MgCl_2_ supplemented with 5 μM Fd and 100 μM NADPH. The periods of actinic- (620 nm, 250 μmol photons m^−2^ s^−1^) and far-red (720 nm, 50 μmol photons m^−2^ s^−1^) illumination are indicated by bright-and dark red bars under the fluorescence data. Periods of darkness are indicated by black bars. ‘SP’ refers to a saturating actinic flash (5000 μmol photons m^−2^ s^−1^). F_o_, F_m_, F_s_ and F_o_′ indicate the fluorescence levels in the dark, during the saturating flash, during the steady state under actinic illumination and under far-red illumination respectively. Data are normalized to F_o_′

The experiments were initiated by measurements of the fluorescence level in the dark, F_0_, followed by a brief pulse saturating actinic light to estimate for the maximal fluorescence yield, F_M_, allowing for normalization of results from preparation to preparation. This treatment was followed by illumination with approximately 250 μmol photons m^−2^ s^−1^ red light for 60 s to establish steady state electron transfer. After illumination, the actinic light was switched off and chlorophyll fluorescence yield were recorded. Upon switching off the actinic light, the fluorescence yield values decayed initially to a level well above the F_0_ value, likely indicating Q_A_ was not completely reoxidized in the dark. To fully oxidize PQ, we periodically illuminated with far-red light (50 μmol photons m^−2^ s^−1^), which preferentially excites PSI, resulted in quenching of the fluorescence signal. Following the far-red light, fluorescence yield increased again, reflecting re-reduction of the PQ pool and Q_A_ by NADPH/Fd. These re-reduction responses, which we term Post-illumination Fluorescence Rise (PIFR) were maintained over multiple cycles of far-red illumination and dark recovery, indicating that the pool of reductant was not depleted during the experiments.

The largest PIFR response was observed in *A. hybridus*, a NAD-ME C_4_ species with an elevated level of NDH in mesophyll chloroplasts compared to bundle-sheath ^33^. The amplitude of the PIFR in *S. oleracea* and *A. thaliana* (Col-0) was approximately 65% and 50% of that observed in *A. hybridus* respectively. The PIFR in Arabidopsis was diminished in the *ndhm* line, which lacks a functional NDH complex ^32^, confirming that the PIFR was related to NDH activity. The residual (20%) PIFR in *ndhm* presumably reflects the activity of the (non-protonmotive) FQR pathway or the recently demonstrated direct reduction of PSII electron acceptors, ^34^. To test these assignments, we repeated experiments in the presence of 1 mM hydroxylamine (HA) and 10 μM DCMU, abolishing variable fluorescence from PSII. The PIFR was inhibited under these conditions (Figure S5), confirming that the PIFR observed in the absence of DCMU and HA was related to PQ pool reduction (mediated by NDH) and subsequent redox equilibration with Q_A_ ^35^.

It is generally considered that the FQR-dependent pathway of CEF is sensitive to AA, whereas the NDH-dependent pathway is insensitive. Addition of 10 μM AA reduced the amplitude of the PIFR by approximately 60% in *S. oleracea* and *A. thaliana* (Col-0) thylakoids, and almost completely in *A. thaliana* (*ndhm*) thylakoid preparations (Figure 4A-D). In contrast, the amplitude of the rise in *A. hybridus* thylakoid preparations was largely unaffected by AA, consistent with an NDH-dominated CEF pathway in this species ^33^. These data provide a useful example for a point that is often overlooked in the literature; that is that both NDH-and FQR activity contribute to the PIFR (which is to be expected, since both activities result in quinone reduction), and the contribution of these activities varies between species. Accordingly, the PIFR is reported to be diminished (but not abolished) in the *A. thaliana pgr5* mutant ^36^, which is considered to be affected in CEF (although see ^37^ for further discussion of this point).

**Figure 4.**
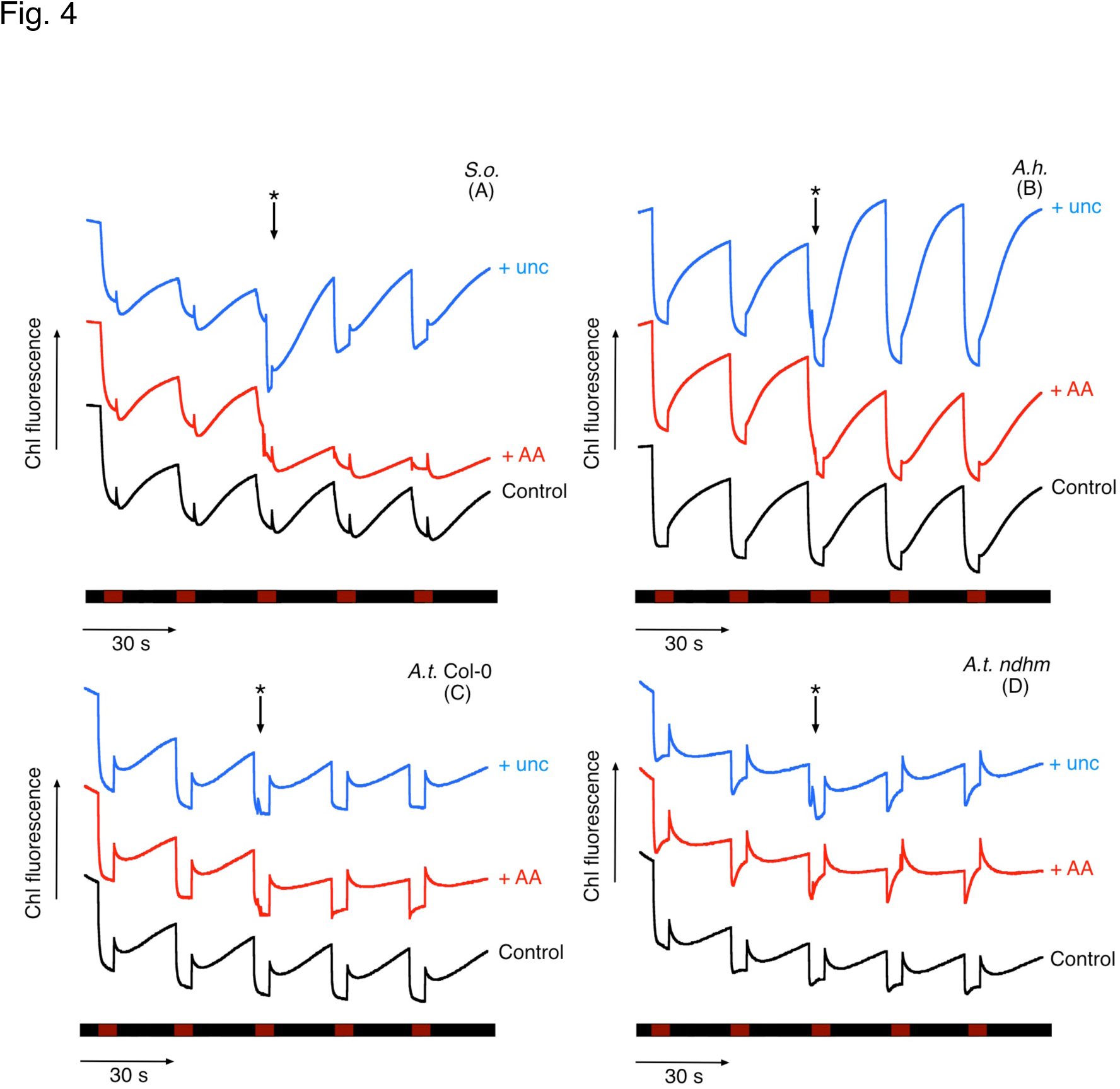
The effect of uncoupling (unc,10 μM nigericin + 10 μM valinomycin) and 10 μM antimycin A (AA) on the post illumination fluorescence rise in (A) *Spinacia oleracea* (*S.o*), (B) *Amaranthus hybridus* (*A.h*.), (C)*Arabidopsis thaliana*- Columbia (*A.t*. Col-0) and (D) *A.t. ndhM* thylakoid preparations. Effectors were added during the course of the experiments as indicated by the asterisked arrow above the fluorescence data. Other conditions as in Figure 3.

Collapsing the transthylakoid Δp through the use of valinomycin and nigericin in the presence of 10 mM K^+^, which should uncouple the thylakoid membranes, resulted in a rapid, two-fold increase in the amplitude and rate of the PIFR observed in *S. oleracea* and *A. hybridus* chloroplast preparations (Figure 4A-D). Dissipation of the Δp allows the protonmotive NDH complex to operate unimpeded by thermodynamic ‘back pressure’ created by the by the light-induced proton gradient across the thylakoid membrane. The PIFR was less sensitive to proton uncoupling in *A. thaliana* (Col-0) and *ndhm* mutant (Figure 4), likely reflecting the larger contribution from the AA-sensitive, and non proton-pumping FQR pathways, which should not be hindered by Δp backpressure.

### Proton-pumping activity of chloroplast NDH probed *in vivo*

*Evidence for proton-pumping associated with the NDH-CEF pathway in vivo*. To assess whether CEF is coupled to proton translocation *in vivo*, we compared the relative changes in light-driven electron and proton transfer reactions between Col-0 and *hcef1*, an Arabidopsis mutant with constitutively elevated rates of CEF through the NDH complex ^38^. Analyses of the results from our previously work [compare Figures 3 and 4 in ^38^] shows that, in *hcef1*, CEF-related increases in light-driven proton fluxes were larger than PSI electron fluxes, suggesting that the H^+^/e^−^ ratio for NDH-related CEF is greater than that for LEF. Given that the H^+^/e^-^ ratio for LEF is expected to be 3, these results suggest that NDH-related CEF has a H^+^/e^-^ ratio of about 4, and thus NDH should pump about 2 protons for each electron passed to the PQ pool. To test this prediction, we compared the increases in light-driven proton flux (*v*_H_^+^) with electron transfer through PSI (*v*_P700_) in wild type and *hcef1* (Figure 5). If NDH does not pump protons, we would expect elevated CEF to decrease the slope of relationship between *v*_H_^+^ and *v*_P700_. If NDH pumped a single proton per electron, the H^+^/e^-^ratio for CEF would be equal to that of LEF, so that the slope of *v*_H_^+^/*v*_P700_ should remain constant with increasing CEF. Finally, if NDH pumped 2 or more H^+^/e^-^ we would expect an increase in *v*_H_^+^/*v*_P700_ with increasing CEF. Our results show a clear increase in *v*_H_^+^/*v*_P700_ in *hcef1* compared to Col-0, supporting the hypothesis that NDH complex pumps more than one (and thus likely two) H^+^/e^-^.

**Figure 5.**
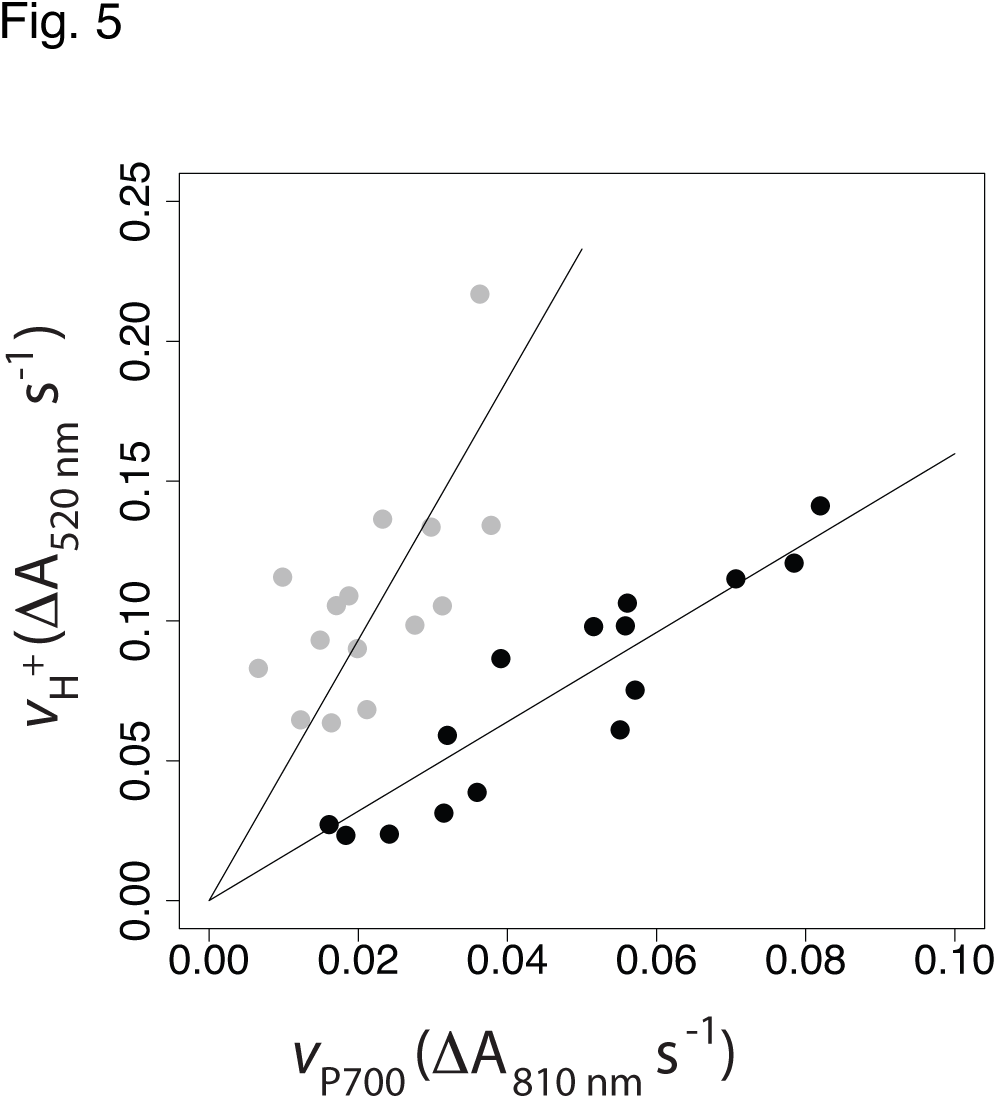
Evidence for involvement of a proton pump in CEF *in vivo*. Relative rates of light-induced thylakoid proton flux (*v*_H_^+^) were estimated by the initial slope of the decay of the electrochromic shift. The rates of turnover PSI (*v*_P700_) were estimated by the rate of decay of the P_700_^+^ absorbance changes upon rapid transition from light to dark, under conditions where P_700_^+^ accumulates (700 nm illumination). Data are from intact leaves of Col-0 (black) and *hcef1* (grey) under increasing light intensities. Lines represent best fit of the data through the origin, with slopes of 1.6 and 4.7 for Col-0 and *hcef1*, respectively.

## Discussion

The function of plastid NDH complex has been the subject of considerable controversy ^39^. NDH is known to catalyze the reduction of PQ by reduced Fd, and thus has often been suggested to play a role in CEF ^18,36,38,40^. However, other CEF pathways, particularly that involving the FQR, have been proposed to constitute the major route for CEF ^14,15,17,41^, though it is clear that CEF can proceed rapidly even in the absence of FQR, through the NDH complex ^36,38,40,42^.

The fact that NDH is homologous to proton-pumping Complex I (NADH:UQ oxidoreductase) of mitochondria and related prokaryotic systems, suggests proton-pumping capacity may also be conserved in plastids, with substantial consequences for the energetics of CEF. In support of this possibility, key residues required for proton-pumping ^22,43,44^ are also conserved in the plastid complex (Figures 1 and S2).

In this work we used several complementary approaches to provide the first direct evidence for the predicted proton-pumping capacity of NDH. In one approach, we developed an *in vitro* assay that reported dark ATP synthesis in coupled, thylakoid preparations under conditions strictly dependent upon NDH activity (Figures 2, S3, S4). Secondly, the extent of the PIFR, a chlorophyll fluorescence phenomenon arising from the NDH-mediated reduction of the PQ pool (Figure 3), increased on addition of uncouplers to thylakoid preparations from *S. oleracea*, *A. hybridus*, and *A. thaliana* (Col-0) (Figure 4). This latter observation is consistent with a protonmotive NDH complex with a pumping (H^+^/e) stoichiometry of 2 or more, so that its enzymatic activity is controlled by the thermodynamic back-pressure from transmembrane Δp. (As discussed below, proton-pumping capacities of less than 2 H^+^/e^-^ should not be thermodynamically limited by physiologically accessible Δp extents.) Finally, *in vivo* data obtained on the high CEF *hcef1* mutant indicated a H^+^/e^-^ stoichiometry for CEF of about 4, requiring an NDH that pumps about two protons per electron transferred to PQ, consistent with the published ratios for respiratory Complex I of about 2H^+^/e^-22,43^.

A proton-pumping NDH should have several important consequences. First, it effectively doubles the amount of ATP produced for a given rate of CEF, making it a more efficient mechanism to balance the ATP/NADPH budget, i.e. compared to non-proton-pumping FQR, fewer turnovers would be required to sustain the required ATP production. The actual contribution of NDH-related CEF to chloroplast bioenergetics has been in dispute. Estimates suggest a chloroplast NDH content of about 0.05 per PSI in *Nicotiana tabacum* ^13^. In order to meet the ATP demands imposed by the CBB cycle, NDH-mediated CEF would need to translocate protons at a rate of ~13% the LEF rate ^45^. With a fixed ratio of 3 H^+^/e^-^for LEF, a 1 H^+^/e^-^ NDH would need to turnover 13% the rate of LEF, while a 2H^+^/e^-^ NDH would need to turnover 9% the rate of LEF. Assuming NDH has an activity similar to that of respiratory Complex I, which has maximal sustained turnover numbers of about 200 e^-^ s^−1^ ^46,47^, we estimate that wild type levels of NDH could support a minimum rate of CEF of about 4 e^-^ s^−1^ per PSI complex, or about 4% of maximal LEF. This rate, in the absence of environmental stresses, is reasonable, as NDH likely works in parallel with the FQR complex and other processes like the malate shunt or the water-water cycle [see review in ^4^] to balance the chloroplast energy budget. Under stressed conditions leading to ROS formation ^48,49^, in chloroplasts of certain C_4_ plants ^33^, and in the high CEF mutant *hcef1* ^38,40^, where high rates of CEF are observed, the content of NDH subunits increase substantially, likely allowing the bulk transfer of electrons through NDH-mediated CEF to be much higher. For example, approximately 15fold higher levels of NDH were observed in the high CEF mutant *hcef1* ^38^.

Furthermore, a proton-pumping NDH is expected to be *required* for non-photosynthetic energy transduction in higher plant plastids, as in the proposed chlororespiratory pathway ^30,50,51^ to involve NDH and the plastid terminal oxidase. Our finding that NDH is a proton pump adds critical support for this possibility because without coupling PQ reduction to proton translocation the proposed pathway would not be able to conserve energy in Δp and ATP synthesis [see also ^26^]

While a proton-pumping NDH provides a higher ATP output for CEF, the coupling of the forward reaction to the generation of Δp should also thermodynamically constrain the extent of the overall reaction. At equilibrium, the relationship between protons translocated into the lumen against Δp per electron falling through a redox span of ΔE_h_ mV is given by equation (1):

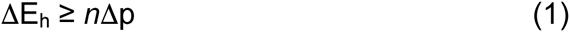

where *n* represents the H^+^/e^-^ ratio. With Fd as electron donor to NDH, ΔE_h_ equals 518 mV assuming 90% reduction of both the PQ and Fd pools (conditions which may be met during photosynthetic induction) at a stromal pH of 7.5. Assuming a Δp of 180 mV across the thylakoid membrane in the light, NDH would be capable of acting as a 2H^+^/e^-^ pump. This assertion also holds for a predominantly oxidized (90%) Fd pool. However, if the redox state of Fd comes into equilibrium with that of NADPH, as would be expected under many conditions (e.g. when electron transfer is limited by turnover of the CBB cycle), then the NDH energetics become significantly more constrained. As illustrated in Figure 6, certain physiologically relevant combinations of redox poise and Δp would limit the forward (plastoquinone reductase) NDH reaction, and even allow for reverse reactions. For instance, when the PQ and NADPH pools become 90% and 50% reduced, respectively, the electron motive driving force for NDH (ΔE_h_) reaches 380 mV, a Δp of 190 mV should limit PQ reduction, and Δp above this should promote the reverse reaction (see below). One may expect these conditions to be produced during rapid light transients where large Δp can be generated ^52^.

**Figure 6.**
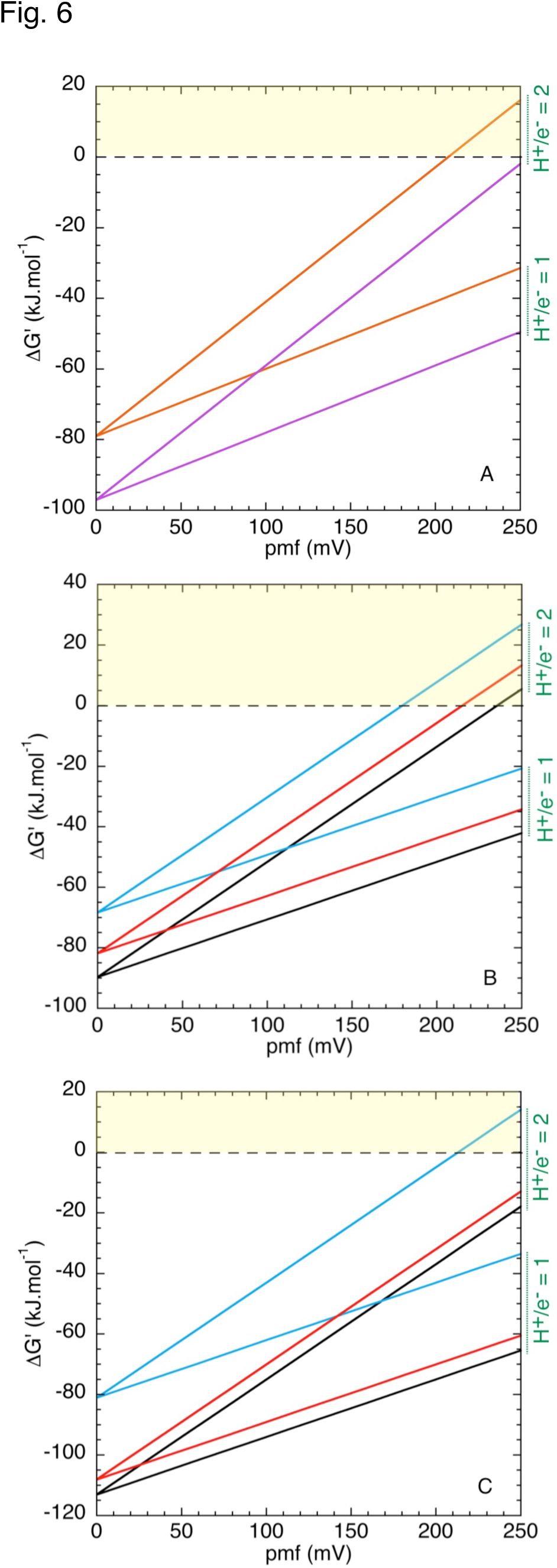
Gibbs free energy plots as a function of protonmotive force (Δp) for the oxidoreductase activity of NDH. Calculations were performed assuming a stromal pH of 7.5 and midpoint potentials (E_m_,_7.5_) of +80, −335 and −430 mV for the PQ/PQH_2_, NADP^+^/NADPH and Fd^3+^/Fd^2+^ couples respectively. T = 298 K. Yellow shaded regions indicate conditions under which the reverse (plastoquinol oxidase) reaction of NDH is energetically favorable. (A) Thermodynamics of NDH-mediated NADPH:PQ oxidoreductase (orange) and Fd:PQ oxidoreductase (purple) activity for H^+^/e^-^ = 1 and H^+^/e^-^ = 2 assuming 50% reduction of donors and acceptors. (B) Thermodynamics of NDH-mediated NADPH:PQ oxidoreductase activity assuming 10% reduced NADPH + 90% reduced PQH_2_ (blue), 90% reduced NADPH + 90% reduced PQH_2_ (red) and 90% reduced NADPH + 10% reduced PQH_2_ (black) for H^+^/e^-^ = 1 and H^+^/e^-^ = 2. (C) Thermodynamics of NDH-mediated Fd:PQ oxidoreductase activity assuming 10% reduced Fd + 90% reduced PQH_2_ (blue), 90% reduced Fd + 90% reduced PQH_2_ (red) and 90% reduced Fd + 10% reduced PQH_2_ (black) for H^+^/e^-^ = 1 and H^+^/e^-^ = 2. Details of the basic thermodynamics equations used to construct this figure are presented in supplemental appendix 1.

The predicted thermodynamic linkage between PQ reduction and proton translocation is clearly demonstrated in our results showing that PIFR associated with CEF is constrained by Δp (Figure 4). It is noteworthy that we would not expect to observe significant thermodynamic backpressure if H^+^/e^-^ was lower than 2. For example, if H^+^/e^-^ for NDH was 1, the equilibrium constant for reduction of PQ by NADPH with Δp=180mV should be 5*10^6^, allowing for essentially full reduction of PQ and Q_A_. This analysis thus strongly supports the involvement of a NDH with a coupling stoichiometry of 2H^+^/e^-^.

The thermodynamic sensitivity of the reaction would provide a simple “self-control” mechanism for regulation of the chloroplast energy budget under conditions of low ATP demand (i.e. high ΔG_ATP_) and favors the partitioning of electrons into downstream metabolism. For instance, a deficit of ATP would favor a low Δp and a more reduced NADPH pool, allowing the NDH reaction to proceed. An excess of ATP might result in high Δp, constraining the reaction, or even force it run in the reverse direction (see below).

The fact that the extent of the NDH reaction is limited by Δp implies that the system reaches quasi-equilibrium. As such, increasing Δp should force the reactions to operate in the “reverse” direction, consuming Δp, and functioning as plastoquinol:NADP^+^ oxidoreductase under conditions of a moderately high Δp ( > 180 mV) when the PQ pool is predominantly (90%) reduced (Figure 6A-C). This condition would fall within the expected ranges of redox and Δp conditions expected in vivo, and might be expected to occur under fluctuating environmental conditions. We note that the proposed reversibility of NDH is not without precedent in nature, and such activity has been observed in prokaryotic homologs of this enzyme ^53,54^.

As can be seen from Figure 6A-C, a 1H^+^/e-coupled NDH would be energetically incapable of significant reverse reaction under physiological conditions. The quinone reductase reaction catalyzed by a 1H^+^/e-NDH (or a non-proton-pumping NDA/NDH2 type complex ^55^) would be insensitive to the presence of uncouplers under physiological conditions and would not manifest the transient post-illumination fluorescence rise response observed in Figure 4 upon valinomycin and nigericin addition. The reverse (NADP^+^ reductase) reaction catalyzed by the 2H^+^/e-thylakoid NDH becomes increasingly energetically favorable as the pool of oxidized acceptor (NADP^+^) and magnitude of Δp increases. With Fd as the electron acceptor, the reversibility of NDH is expected to be strongly limited below a Δp of ≈ 200 mV (Figure 7C). However, rapid oxidation of Fd molecules reduced by NDH activity would facilitate this reverse reaction.

Thus, under conditions of high Δp, we propose that chloroplasts may conduct electron flow from water, via PSII, to NADPH without the direct participation of PSI, effectively a ‘pseudo-linear’ pathway for photosynthetic electron transfer that could operate under special conditions of high Δp and reduced PQH2, helping to balance the ATP/NADPH energy budget For example, activation of the pseudo-linear pathway may have large consequences for regulation of metabolic processes that are modulated by redox state of NADPH through the thioredoxin system, possibly explaining why NDH knockouts exhibit strong phenotypes during the induction of photosynthesis, when high Δp and imbalances of redox state are expected, or when subjected to environmental stress, such as decreased CO2, fluctuating light intensity or chilling ^56,57^.

The trade-offs between efficiency and thermodynamic constraints may also explain why there are multiple routes of differentially-regulated CEF pathways in chloroplasts. Under some conditions, highly efficient, but slower, synthesis of ATP via the NDH complex would be favored. However, NDH activation may be slow, taking approximately 20 minutes ^40^, and a more rapidly-activated pathway may be needed for rapid responses. In addition, there may also be conditions where higher Δp than that afforded by the NDH pathway may be beneficial, perhaps to acidify the lumen and down-regulate photosynthesis ^9^. We thus propose that the NDH and FQR modes of CEF are activated step-wise when the downstream metabolism decreases ATP/NADPH. First, a reducing environment is generated as NADPH accumulates, and CEF activated through the non-proton-pumping, FQR-dependent (antimycin A-sensitive) pathway. Under conditions in which this route of CEF is not able to augment the ATP deficit, and restore redox homeostasis to the chloroplast, ROS is generated and activates the proton-pumping NDH complex ^40,48,49^. This would increase the yield of ATP formation per e^-^ transfer, increasing the efficiency of ATP production via CEF. This route of CEF is not as rapidly activated as the FQR, and long term ROS generation leads to not only activation of already assembled complexes, but an increase in total NDH content ^40,48,49^.

## Methods

### Reagents, plant materials and growth conditions

Reagents were purchased from Sigma-Aldrich (St Louis, MO) unless otherwise noted.

*Arabidopsis thaliana* and *Amaranthus hybridus* (Swallowtail Garden Seeds, Santa Rosa, CA) plants were grown in a 16:8 light:dark photoperiod. Experiments using Col-0 and were performed at 3-4 weeks of age, while slow-growing *hcef1* was used at the same developmental stage as Col-0 (around 6-7 weeks of age). Chloroplasts were prepared from *A. hybridus* Col-0 and *ndhm* (Salk_087707) plants 3 weeks after germination. Spinach was obtained from a local market, with chloroplasts prepared on the morning of purchase.

Chloroplasts were prepared from *S. oleracea* and *A. hybridus* leaves as described in ^58^, and chlorophyll determined as in ^59^.

### Spectroscopic measurements

All spectroscopic measurements were made using an integrated diode emitter array (IDEA) spectrophotometer/fluorimeter ^60^. Plants were poised by illuminating with 700 nm actinic light to favor oxidation of PSI and enable measurement of electron flux through PS I by analysis of dark interval relaxation kinetics.

Transthylakoid proton flux (*v*_H_^+^, Δ_A520 nm_ m^−2^ s^−1^) was calculated using the electrochromic shift of the carotenoids at 520 nm (as described in ^40,61^, Figure S6A). To correct for variability in leaf pigmentation between Col-0 and *hcef1* total extent of ECS in *hcef1* was normalized to the fraction of Col-0 per area leaf chlorophyll content as described previously for *hcef1* ^38,42^.

Redox state of PSI was monitored using absorbance changes at 820 nm in a protocol modified from ^62^. The initial rate of P700^+^ re-reduction kinetics were used as the relative rate of electron transfer through PSI (*v*_P700_, ΔA_820 nm_ m^−2^ s^−1^) (Figure S6B).

Postillumination chlorophyll *a* fluorescence transients were measured as described in ^13,31^ with modifications. Thylakoids were preilluminated with 250 μmoles photons m^−2^ s^−1^ (620 nm), for 1 minute. After the light to dark transition, the measuring beam (505 nm, 0.5 μmol photons m^−2^ s^−1^) was pulsed at a 10 Hz interval. Chlorophyll fluorescence changes during the dark interval were monitored for 145 seconds, with 5 second pulses of far-red illumination (720 nm, 50 μmol photons m^−2^ s^−1^) every 20 seconds.

The H^+^/e^-^ of CEF in *hcef1* was estimated by the following calculation:

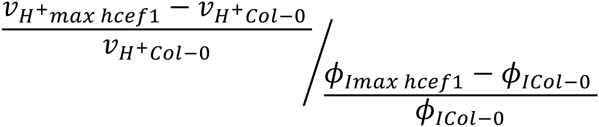

in which the Col-0 values correspond with a constant LEF.

### In vitro *ATP synthesis assays*

Proton-pumping *in vitro* was measured as ATP production in the dark using the Promega (‘Enliten’) luciferase/luciferin reagent kit with a laboratory-constructed photomultiplier-based phosphoroscope. Osmotically ruptured *S. oleracea* chloroplasts (prepared in the presence of 2 mM reduced DTT) were present at a chlorophyll concentration of 50 μg/ml. DCMU was present at 10 μM. Fd and NADPH were present at 5 μM and 100 μM respectively. The assay buffer (pH 7.5) consisted of 10 mM HEPES, 2 mM potassium phosphate, 10 mM KCl, 5 mM MgCl_2_, 2 mM DTT and was supplemented with 2 mM ADP and 100 μM diadenosinepentaphosphate (an adenylate kinase inhibitor). Premixed ‘Enliten’ recombinant luciferase/luciferin reagent (used as supplied by Promega) was added to a final concentration of 8% (v/v). Proton-pumping was initiated by the addition of 50 μM decylplastoquinone (dPQ), and the proton gradient was collapsed by the addition of 10 μM nigericin and 10 μM valinomycin. The plastid ATP synthase was activated by red actinic illumination (50 μmoles photons m^−2^ m^−1^, 625 nm) of the chloroplast suspension for two minutes immediately prior to the addition of DCMU, ADP, ‘Enliten’ reagent and the start of data collection.

## Acknowledgements

The authors would like to thank Dr. Jeffrey Cruz for helpful discussions. This work was supported by Grant DE-FG02-11ER16220 from the Photosynthetic Systems program from Division of Chemical Sciences, Geosciences, and Biosciences, Office of Basic Energy Sciences of the US Department of Energy (to DMK). We dedicate this paper to the memory of Dr. Derek Bendall (1930-2014), pioneer of cyclic electron flow research and mentor to N.F.

